# Using machine learning to understand microgeographic determinants of the Zika vector, *Aedes aegypti*

**DOI:** 10.1101/2022.03.03.482880

**Authors:** Jagger Alexander, André Barreto Bruno Wilke, Alejandro Mantero, Chalmers Vasquez, William Petrie, Naresh Kumar, John Beier

## Abstract

There are limited data on why the 2016 Zika outbreak in Miami-Dade County, Florida was confined to certain neighborhoods. In this research, *Aedes aegypti*, the primary vector of Zika virus, are studied to examine neighborhood-level differences in their population dynamics and underlying processes. Weekly mosquito data were acquired from the Miami-Dade County Mosquito Control Division from 2016 to 2020 from 172 traps deployed around Miami-Dade County. Using Random Forest, a machine learning method, predictive models of spatiotemporal dynamics of *Ae. aegypti* in response to meteorological conditions and neighborhood-specific socio-demographic and physical characteristics, such as land-use and land-cover type and income level, were created. The study area was divided into two groups: areas affected by local transmission of Zika during the 2016 outbreak and unaffected areas. *Aedes aegypti* populations in areas affected by Zika were more strongly influenced by 14- and 21-day lagged weather conditions. In the unaffected areas, mosquito populations were more strongly influenced by land-use and day-of-collection weather conditions. There are neighborhood-scale differences in *Ae. aegypti* population dynamics. These differences in turn influence vector-borne disease diffusion in a region. These results have implications for vector control experts to lead neighborhood-specific vector control strategies and for epidemiologists to guide vector-borne disease risk preparations, especially for containing the spread of vector-borne disease in response to ongoing climate change.

## Introduction

Vector mosquito populations in Miami-Dade County, Florida pose a great threat to the local population health as well as the United States at large, as Miami-Dade may serve as an entry point for vector-borne diseases [1]. In the most recent case, in June of 2016, Miami-Dade was the site of the first confirmed Zika virus disease cases in the United States [2, 3]. *Aedes* mosquitoes are the primary vectors of the Zika virus. *Aedes aegypti* specifically is the most important mosquito vector in the transmission of the Zika virus in the United States [4]. In South Florida, across the United States, and over the planet, vector-borne diseases have the potential to spread at increased rates and over broader distributions due to climate change [5]. Understanding the patterns that control vector species’ behavior and population dynamics is critical to mosquito control efforts and would be beneficial to public health.

Variations in vector mosquito abundance and vector-borne disease spread with weather patterns have been modeled for decades [6-12]. Studies across different continents and habitat types show significant correlations between vector mosquito abundance and temperature, humidity, and precipitation. Though some mechanisms for these variations remain unresolved, some proposed mechanisms include different activity levels and reproductive rates in different temperatures and humidities, different availability of aquatic larval habitats with precipitation, and changing interspecies interactions with weather conditions [1].

Species abundance also changes with habitat, as different habitats offer different ecology, resource availability, and larval habitat potential. Urban environments have unique properties that can often support vector species and lead them to a greater interaction with humans [13-15]. In Miami-Dade County, ongoing research has shown that features of the urban environment such as tire shops [16], cemeteries [17], urban farms [18], and ornamental bromeliads [19] all provide aquatic habitats for mosquito development [20]. The relationships of these vector species to their urban environment are dependent on weather and climate; water is a determining factor in the physical environment and is necessary for them to breed, while temperature inherently affects mosquito energy expenditure, activity, and physiology [21]. Thus, while the urban environment has been studied extensively as a habitat for vector mosquitoes, it is important to understand how weather patterns interact with various types of urban habitats, such as those where buildings are denser or those that offer more vegetation, and in turn influence mosquito abundance.

As urban habitats represent an important intersection between vector species and human society, it is important that we understand the population dynamics of vector species populations in these environments. The motivation for this analysis is drawn from unique variations across Miami-Dade County in the spread of the Zika virus. Local transmission was generally confined to certain regions across the county and was stopped after mosquito control through the implementation of aerosol insecticides [2, 3]. According to our best knowledge, there are currently few studies examining why the 2016 Zika virus outbreak in South Florida was concentrated in select areas.

Here, long-term trends are analyzed separately in *Ae. aegypti* populations in regions affected by Zika during the 2016 outbreak and in regions that were unaffected. It is hypothesized that different regions across Miami-Dade County, particularly those that sustained local transmission of Zika virus and those that did not, will have different population trends (differences in the mean and variance of *Ae. aegypti* as well as differences in observed temporal patterns). It is also hypothesized that weather variables have differing relationships with *Ae. aegypti* populations in Zika affected regions as opposed to *Ae. aegypti* populations in unaffected regions.

To test these hypotheses, a machine learning approach was used. Random Forest, a supervised learning method, can be used for multivariate regression analyses and determining the relative importance of multiple predictor variables on an outcome [22]. This approach has rarely been used for vector-species population and vector-disease modeling [23, 24] but has been used extensively in other fields, with some extensions to the method having been specifically implemented for spatial analysis [25]. This novel method may provide a better approach to understanding the complexity of *Ae. aegypti* population drivers in urban habitats.

## Material and Methods

BG-Sentinel traps (BioGents, Regensburg, Germany), baited with CO_2_, were utilized around Miami-Dade County to collect data on *Ae. aegypti* populations [26]. Around the county, 172 traps were placed in total, with 54 placed in the areas sustaining local transmission of Zika during the 2016 outbreak and 118 placed in the unaffected region, or non-Zika region. Data were collected weekly over a period from July 2016 to March 2020. Fieldwork and data collection were done by the Miami-Dade County Mosquito Control. Data is only included after 2016 when collection became more intensive in response to the Zika virus outbreak. As such, mosquito traps were placed in areas affected by Zika about a month before they were placed in unaffected areas. Additionally, the start dates and end dates of each trap’s implementation varied. A full map of trap locations is included in the Results section (Fig 1). While trap locations were limited to areas where volunteers were willing to keep the trap on their properties, active traps were consistently placed at least 50 meters apart. According to the manufacturer, each trap has a maximum collection range of 20 meters, so the minimum of 50 meters distance is sufficient to prevent overlap between collections and keep sampling power constant [27]. Traps were deployed weekly for 24-hour periods. Mosquitoes were sexed and identified by species by Miami-Dade County Mosquito Control. As the traps were baited with CO_2_ to attract adult female *Ae. aegypti* searching for a blood meal, any male mosquitoes in the traps were considered coincidental and uninformative for surveillance purposes. They were therefore excluded in the data used for this study.

**Fig 1.**
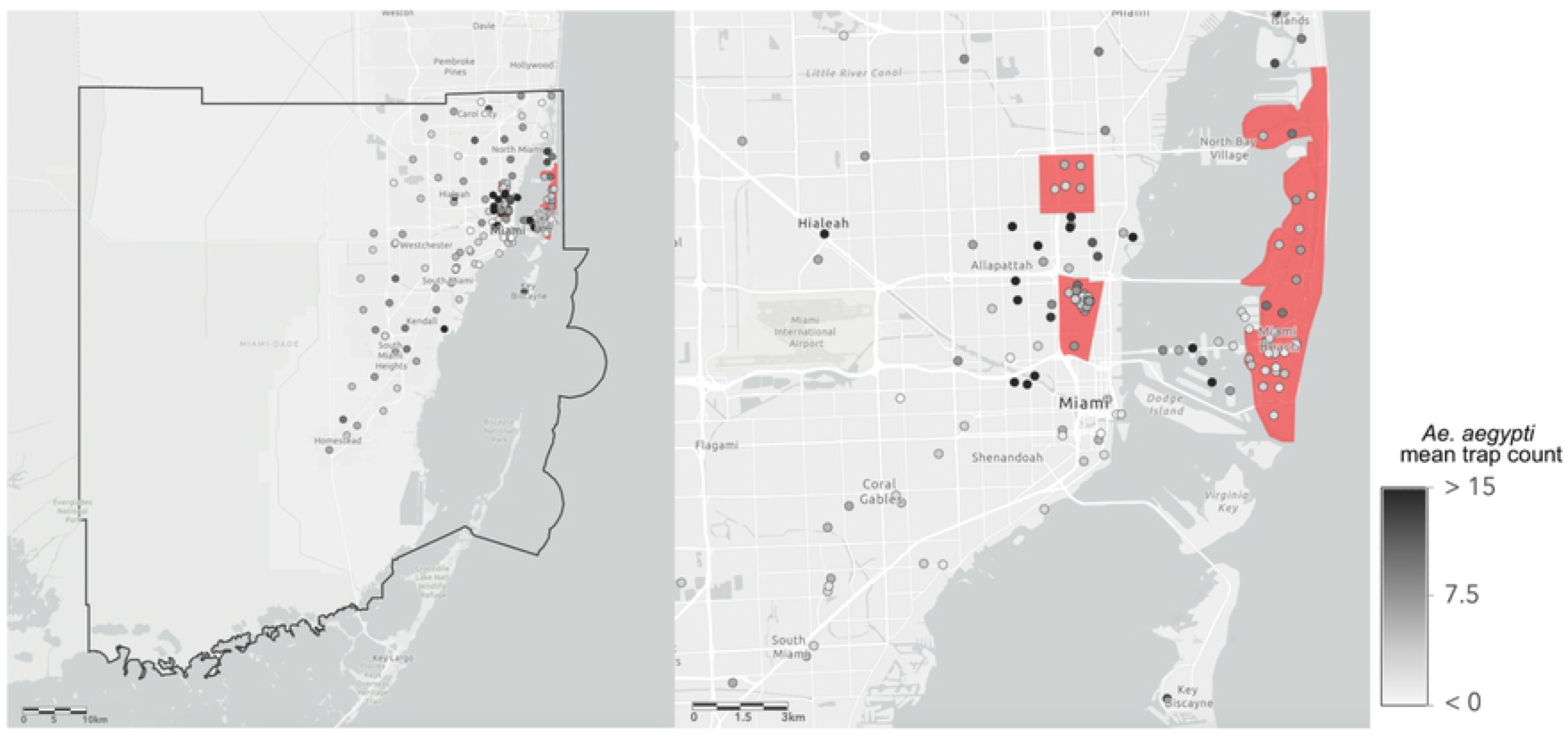
Map of Miami-Dade County Mosquito Control traps. The map on the left includes all 172 traps from which data was collected for this study. The black bold line represents the boundary of Miami-Dade County. The map on the right highlights areas (shaded red) that sustained local Zika virus transmission during the 2016 outbreak. Trap locations are shaded according to the mean value of *Aedes aegypti* collected by that trap over each collection during the study period (2016 – 2020).

Daily high temperature, daily low temperature, and daily total precipitation data over the period from June 2016 to March 2020 were gathered from National Oceanic and Atmospheric Administration Online Weather Data, as accessed in May 2020 [28]. Daily average wind speed and relative humidity were gathered from the National Centers for Environmental Information Local Climatological Data as accessed in May 2020 [29]. The original collection site of each of these data was Miami International Airport, Station USW00012839, which is located at 25.7881° N, 80.3169° E.

Land-use data was accessed online from Miami-Dade County Open Data Hub [30]. Three categorical variables were used to describe land use in this study. The first was major land use or a description of the site where the trap was placed. This variable had 11 values: ‘Low-Density Residential (LD),’ ‘Medium-Density Residential (MD),’ ‘High-Density Residential (HD),’ ‘Business,’ ‘Education,’ ‘Religious,’ ‘Attraction,’ ‘Hospital,’ ‘Industrial,’ ‘Park,’ and ‘Vacant.’ The other two categorical land-use variables are minor land-use A and minor land-use B. To acquire these, the two dominant land-use types within a 100-meter radius of the original trap site were categorized. This process was done using ArcGIS Pro version 10.8.1 [31]. All of the categories of the major land-use variable remain the same for major land-use A and major land-use B except for the addition of ‘Road,’ ‘Lake,’ ‘Rail,’ and ‘Bay.’ In the case that all the land within 100 meters of the trap was categorized as only one type, that one type was recorded for major land-use and both minor land-uses. In the case that all land surrounding the trap (within 100 meters) was categorized as only one type, both minor land-uses were recorded as the same. Per capita income data was acquired from the American Community Survey (2015 – 2019) [32] and is provided by the census tract. These data were accessed online through a third-party interface [33] and verified for accuracy against the original census data.

Each trap is sorted as belonging to one of two regions: areas that were affected by local transmission of Zika virus during the 2016 outbreak (“Zika region”), and areas that were not affected by local transmission of Zika virus during the 2016 outbreak (“non-Zika region”). The exact boundaries of these locations are shown in the results. Further, data are grouped into six sub-regions; three areas affected by local Zika virus transmission (Little River, Miami Beach, Wynwood), and three unaffected (Downtown/Brickell, Hialeah, Homestead). Trap counts from each region and sub-region were fitted to a Poisson distribution and standard deviation and λ statistics were calculated, including a 95% confidence interval for λ. This analysis was conducted using the MASS package 7.3-54 [34] in R version 4.11 [35] run through R studio version 1.4.1717 [36].

A time series plot was produced by aggregating all trap counts in the Zika and non-Zika regions to the weekly level and then creating a three-week moving average curve of the data in Microsoft Excel for Mac version 16.54 [37]. A temporal autocorrelation plot was produced for the weekly aggregated data using the stats package version 3.6.2, one of the core packages in R [35].

A data frame was constructed by co-locating the *Ae. aegypti* adult female counts collected at each individual trap collection with appropriate spatial variables (major land-use, minor land-use A, minor land-use B, and per capita income), as well as temporal variables (average wind speed, daily maximum temperature, daily minimum temperature, relative humidity, precipitation) on the day of collection, one day before collection, seven days before collection, fourteen days before collection, and twenty-one days collection. The choice of these lag times is intentional to provide weather conditions over the adult life and breeding cycle of this species. Lag times in between 0, 1, 7, 14, and 21 days are avoided as excess autocorrelation between variables would reduce the interpretability of Random Forest models. A table of all explanatory variables included in analyses is shown below (Table 1).

**Table 1.** Variable names, abbreviations, and sources for each explanatory variable. Each of these variables is used in random forest predictive models against *Aedes aegypti* trap counts. Numbers next to an abbreviation correspond to lag times used (days before collection). Further explanations of each variable are provided in methods.

| Variable | Abbreviation(s) | Source | Spatial/Temporal |
| --- | --- | --- | --- |
| Average daily wind speed | AWND (0, 1, 7, 14, 21) | NCEI Local Climatological Data [29] | Temporal |
| Daily maximum temperature | TMAX (0, 1, 7, 14, 21) | NOAA NOW Data [28] | Temporal |
| Daily minimum temperature | TMIN (0, 1, 7, 14, 21) | NOAA NOW Data | Temporal |
| Average daily relative humidity | RH (0, 1, 7, 14, 21) | NCEI Local Climatological Data | Temporal |
| Total daily precipitation | PRCP (0, 1, 7, 14, 21) | NOAA NOW Data | Temporal |
| Major land-use | LUM | Miami-Dade County Open Data Hub [30] | Spatial |
| Minor land-use A | LUmA | Miami-Dade<br>County Open Data<br>Hub | Spatial |
| Minor land-use B | LUmB | Miami-Dade<br>County Open Data<br>Hub | Spatial |
| Per Capita Income | PCI | US Census Bureau –<br>American<br>Community Survey<br>[32] | Spatial |

Random Forest analyses were performed using the RandomForestSRC package version 2.12.1 in R [38]. Three random forest models were created: one for all trap counts, a second for only those in the Zika regions, and a third for only those in the non-Zika regions. For each model, all explanatory variables listed in the data frame above were used as predictive values. Individual trap, region, sub-region, and date were not used as predictive values as the inclusion of any of these values would detract from understanding the effects of the selected weather and spatial variables on mosquito counts across each region. Variable importance (VIMP) scores shown are normalized by running two new Random Forest models over the same predictive variables but instead against the z-score of Aedes aegypti trap counts. These separate z-score based Random Forest models are only used for VIMP score calculations.

To compare the differences between the random forest models generated for Zika and non-Zika regions, cross-prediction was performed between the models. First, the weather and spatial data from the non-Zika region was inputted into the random forest model generated for the Zika region to predict the non-Zika region’s *Ae. aegypti* trap counts. This is plotted on the same axis as the predictions of the non-Zika model for the non-Zika trap counts. Similarly, the explanatory data from the Zika region was inputted into the Random Forest model generated for the non-Zika region to predict the Zika region’s *Ae. aegypti* trap counts; this is plotted on the same axis as the predictions from the Zika model. Using base R, root-mean-square error (RMSE) was calculated for each prediction to compare their strength. Finally, test scenarios are presented to demonstrate the different predictive outputs of each model. These test scenarios are chosen to ensure they are within the appropriate sample space of the model. Input conditions are included as supporting information.

## Results

In Zika and non-Zika regions, 118 and 54 traps were placed respectively (Table 2). In total, in the Zika and non-Zika regions, there were 15,213 and 11,154 collections respectively. Trap counts show that there is spatial (Fig 1) and temporal variation in *Ae. aegypti* across Miami-Dade County (Fig 2). The 95% confidence intervals indicate that the λ statistic for Zika regions was significantly lower than non-Zika regions (p < 0.05) (Table 2). The λ statistics were significantly different across all sub-regions except for Miami Beach and Little River (p < 0.05).

**Table 2.**
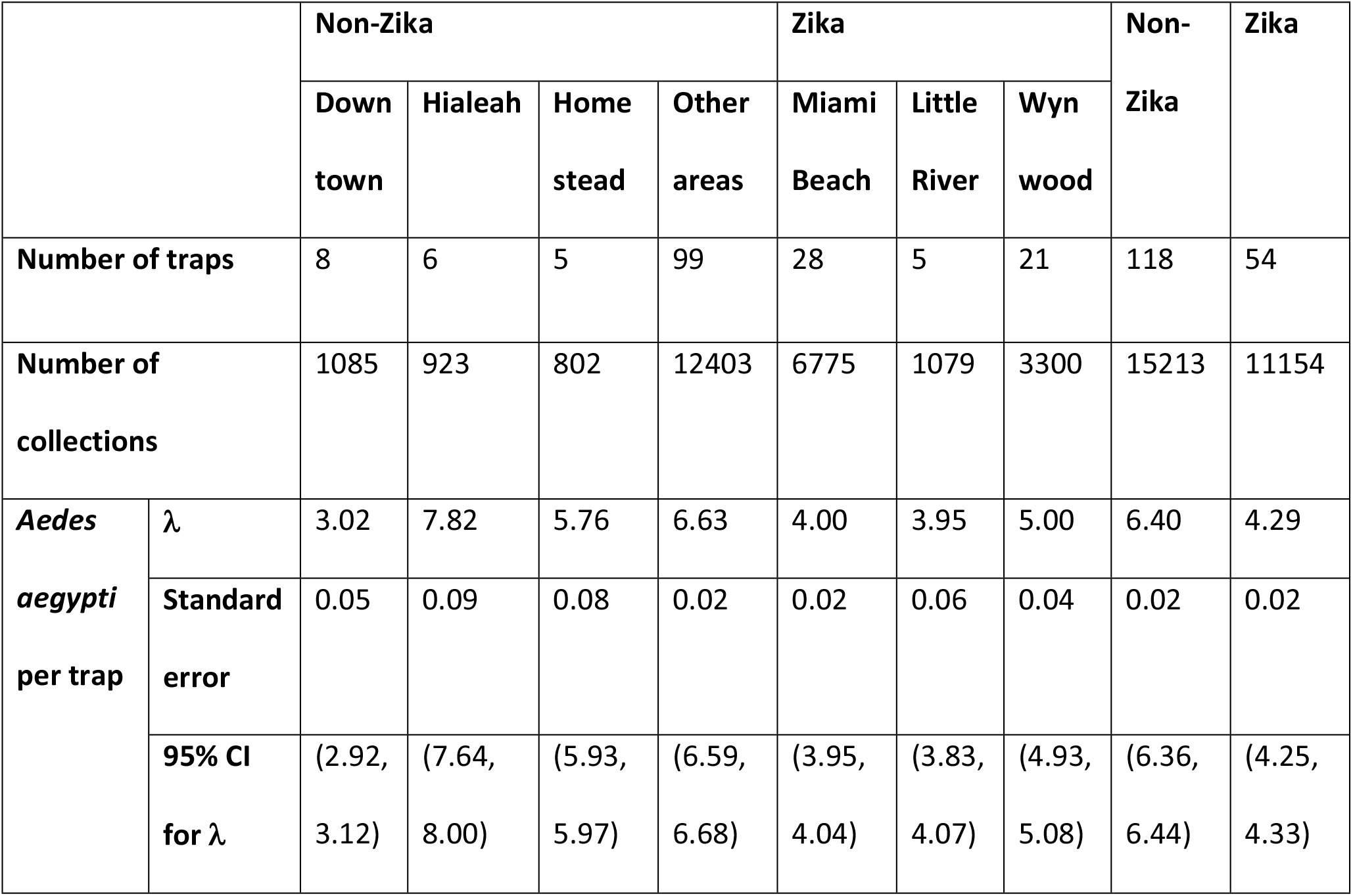
Summary statistics for *Aedes aegypti* collected at areas across Miami-Dade County. Data are grouped into locations affected by local Zika virus transmission during the 2016 outbreak and unaffected. Areas are further subdivided into six defined regions. Trap counts from each region were fitted to a Poisson distribution. Appropriate statistics and 95% confidence intervals for λ are shown.

**Fig 2.**
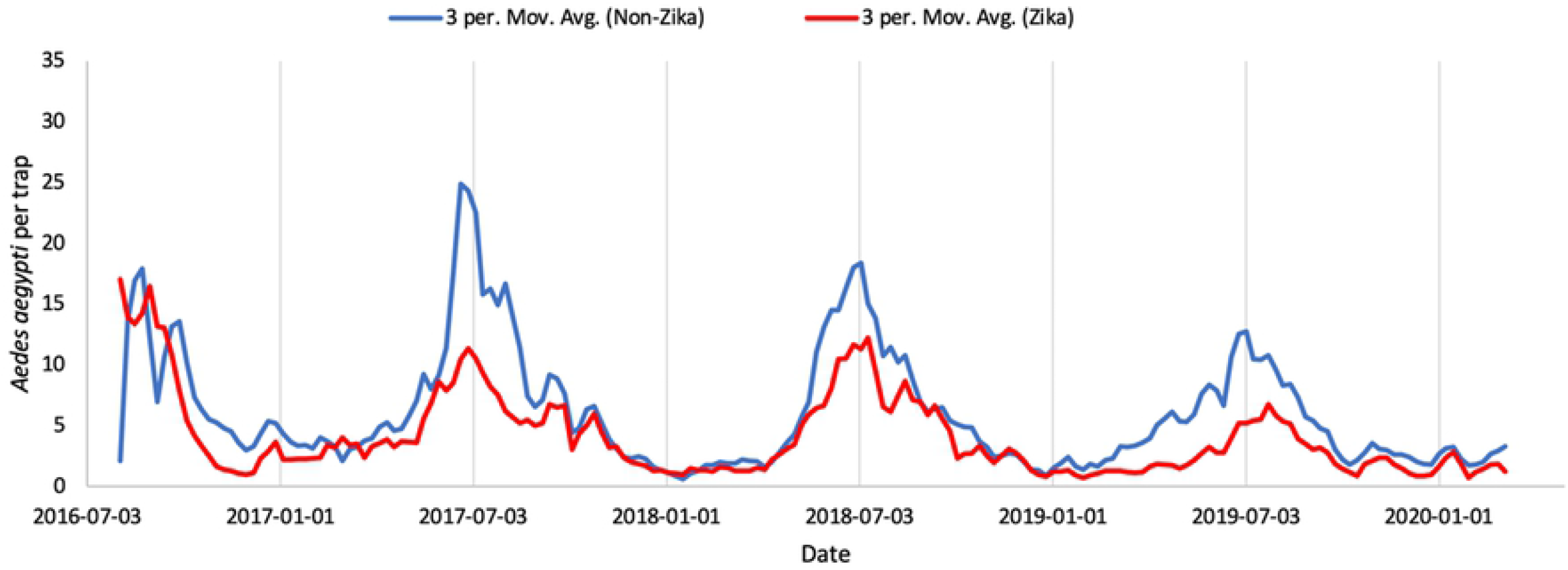
3-period moving average curve of *Aedes aegypti* per trap. Conducted for daily average of all traps areas affected by local Zika transmission during the 2016 outbreak (red) and unaffected areas (blue).

*Aedes aegypti* populations fluctuate over the year broadly and from week-to-week (Fig 2). The populations are highest in both regions in the summer months, peaking around July, and lower from mid-fall through mid-spring. *Aedes aegypti* populations in Zika regions are generally lower than populations in non-Zika regions over the summer months though the difference between the two is smaller in the winter months. Populations in both regions, especially during peak season, decreased slightly over the study period from 2016 to 2020. In Zika regions there is significant temporal autocorrelation between weekly-aggregated trap counts for up to 25 weeks, while for non-Zika regions there is significant temporal autocorrelation between weekly-aggregated trap counts for over 40 weeks (Fig 3). In both cases, this difference represents a period of time greater than the noticeable seasonal differences (Fig 2).

**Fig 3.**
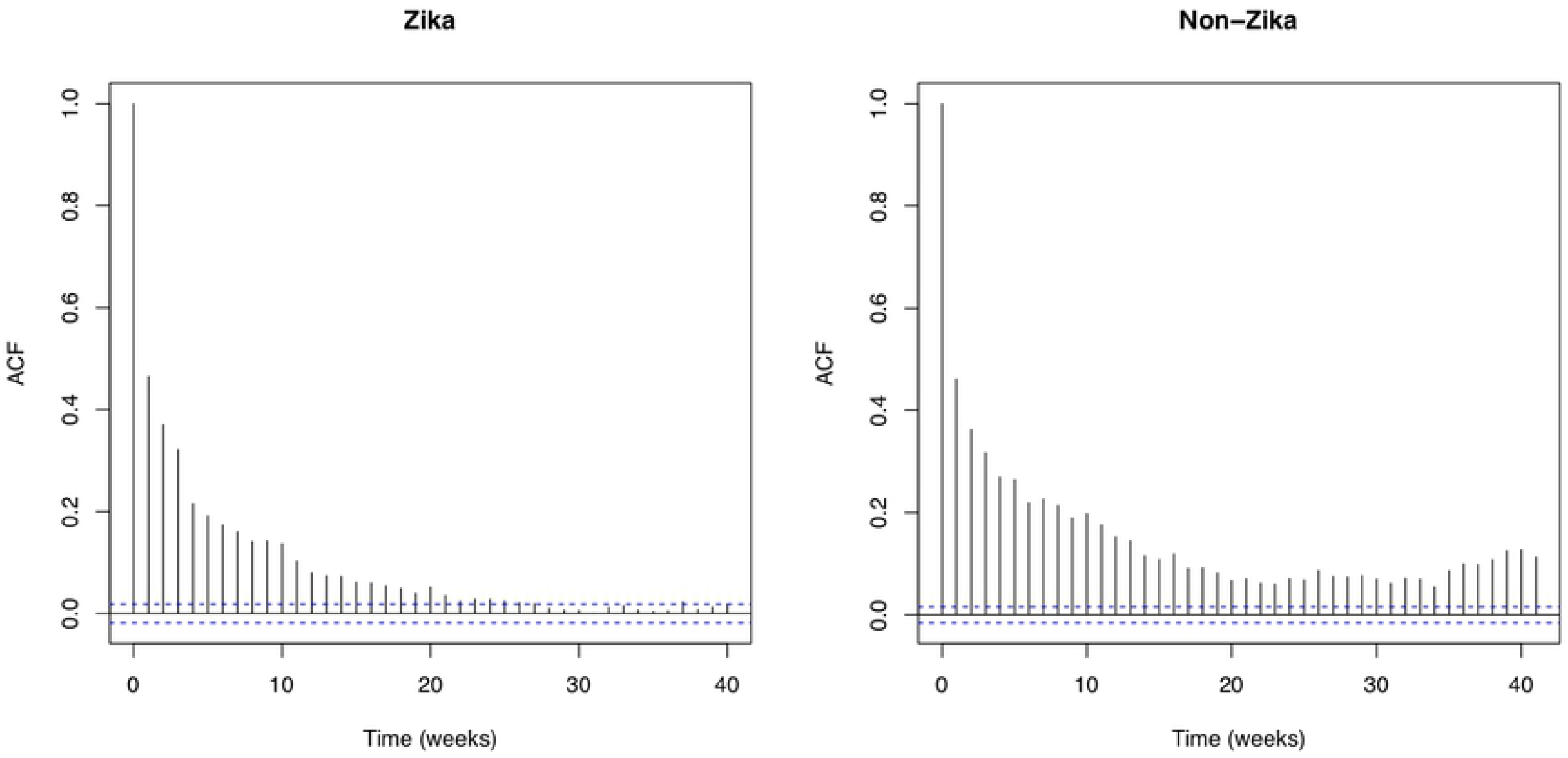
Temporal autocorrelation plot of average mosquito counts. These plots are shown over any one collection date for all traps in areas affected by Zika during the 2016 outbreak (left) and unaffected areas (right). Values outside of the blue-dashed lines are significantly different from zero.

Random Forest model output for Zika and non-Zika regions indicates that the model explains about 40% of the variation in *Ae. aegypti* counts in the Zika region, 9% more than the model for the non-Zika region, and about 10% more than a model of all data across the county (Table 3). Variable importance scores show that per capita income and maximum daily temperature on the day of collection and the day before collection were the most important predictors for both models (Fig 4). For the Zika model this is followed by maximum daily temperature 14 days before collection, minimum daily temperature the day of collection, relative humidity fourteen days before collection, minimum daily temperature 14 days before collection, and precipitation 21 days before collection, whereas for the non-Zika model this is followed by major land-use and minor land-use A.

**Table 3.** Details of Random Forest models. Models were trained on data of adult female *Aedes aegypti* counts from traps in areas affected by local transmission during the 2016 Zika outbreak, traps in unaffected regions, and all traps.

|  | **Zika** | **Non-Zika** | **All** |
| --- | --- | --- | --- |
| **N (number of trap collections)** | 11154 | 15213 | 26367 |
| **Average number of terminal nodes** | 1290.281 | 1876.577 | 3198.846 |
| **Resample size used to grow trees** | 7049 | 9615 | 16664 |
| **Number of trees** | 1000 |  |  |
| **Forest terminal node size** | 5 |  |  |
| **Variables used** | Maximum temperature, minimum temperature, precipitation, relative humidity, wind speed (each at 0, 1, 7, 14, and 21 day lag times), major land-use, minor land-use A, minor land-use B, per capita income |  |  |
| **R<sup>2</sup>** | 0.403 | 0.311 | 0.3042605 |

**Fig 4.**
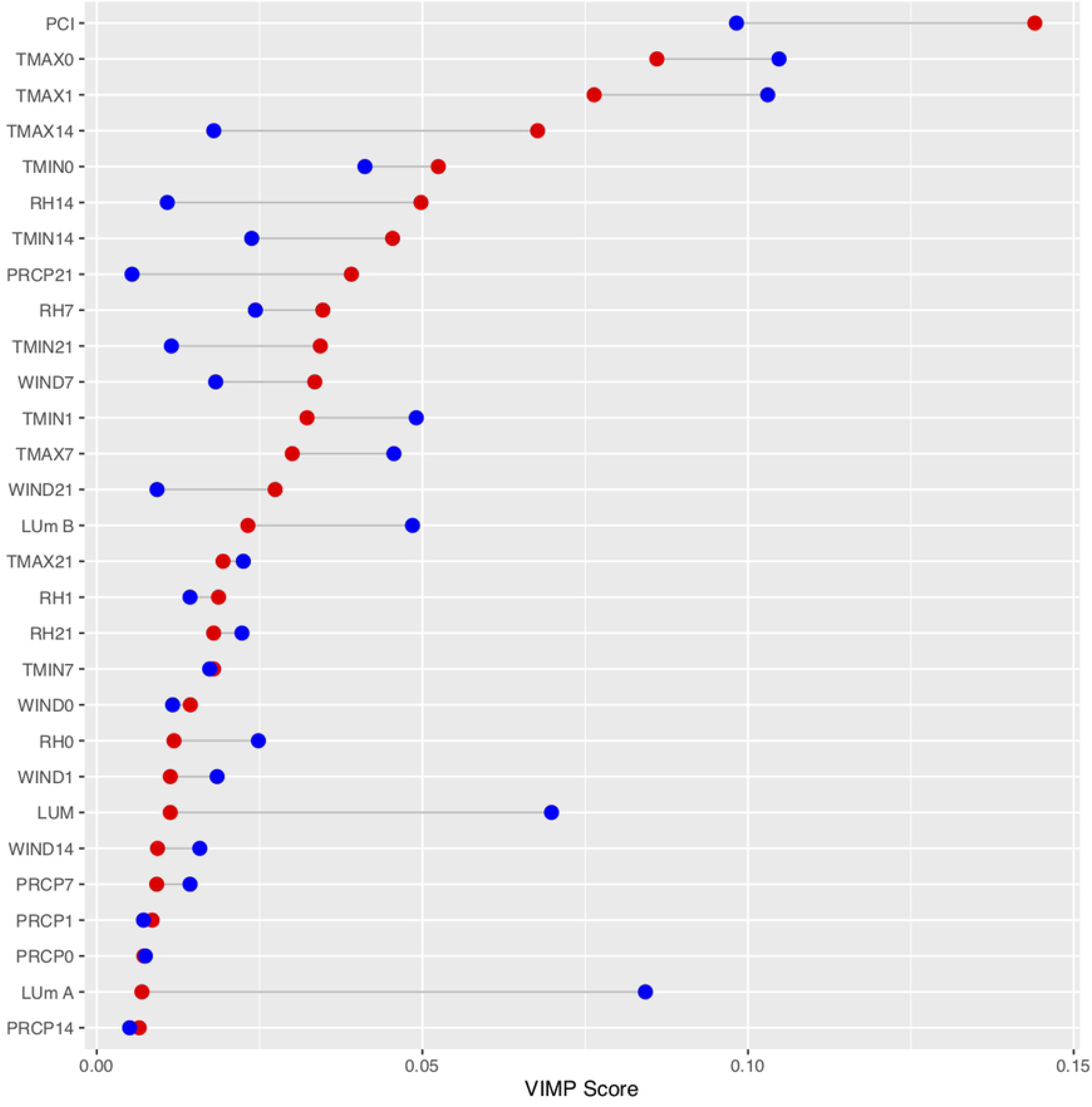
Variable importance (VIMP) scores for Random Forest runs. These scores are shown for models of *Aedes aegypti* adult females collected in traps in Zika affected regions of Miami (red) and areas of Miami unaffected by the Zika outbreak (blue). VIMP scores are “normalized” by modeling against the z-score of *Ae. aegypti* collected. Numbers next to variables indicate associated lag times.

Partial plots of variables in the Random Forest model show the partial effect of each variable towards the model (Fig 5). Results indicate similar trends in maximum and minimum temperature at 14-day lag times as they contribute to expected *Ae. aegypti*. For relative humidity 14 days before collection and precipitation 21 days before collection, increases in either has a greater effect on the expected *Ae. aegypti* value in the Zika regions than the non-Zika regions. For per capita income in both models, at low incomes there is a constant expected *Ae. aegypti* count, a dip in expected *Ae. aegypti* at medium incomes, and higher counts in higher income areas.

**Fig 5.**
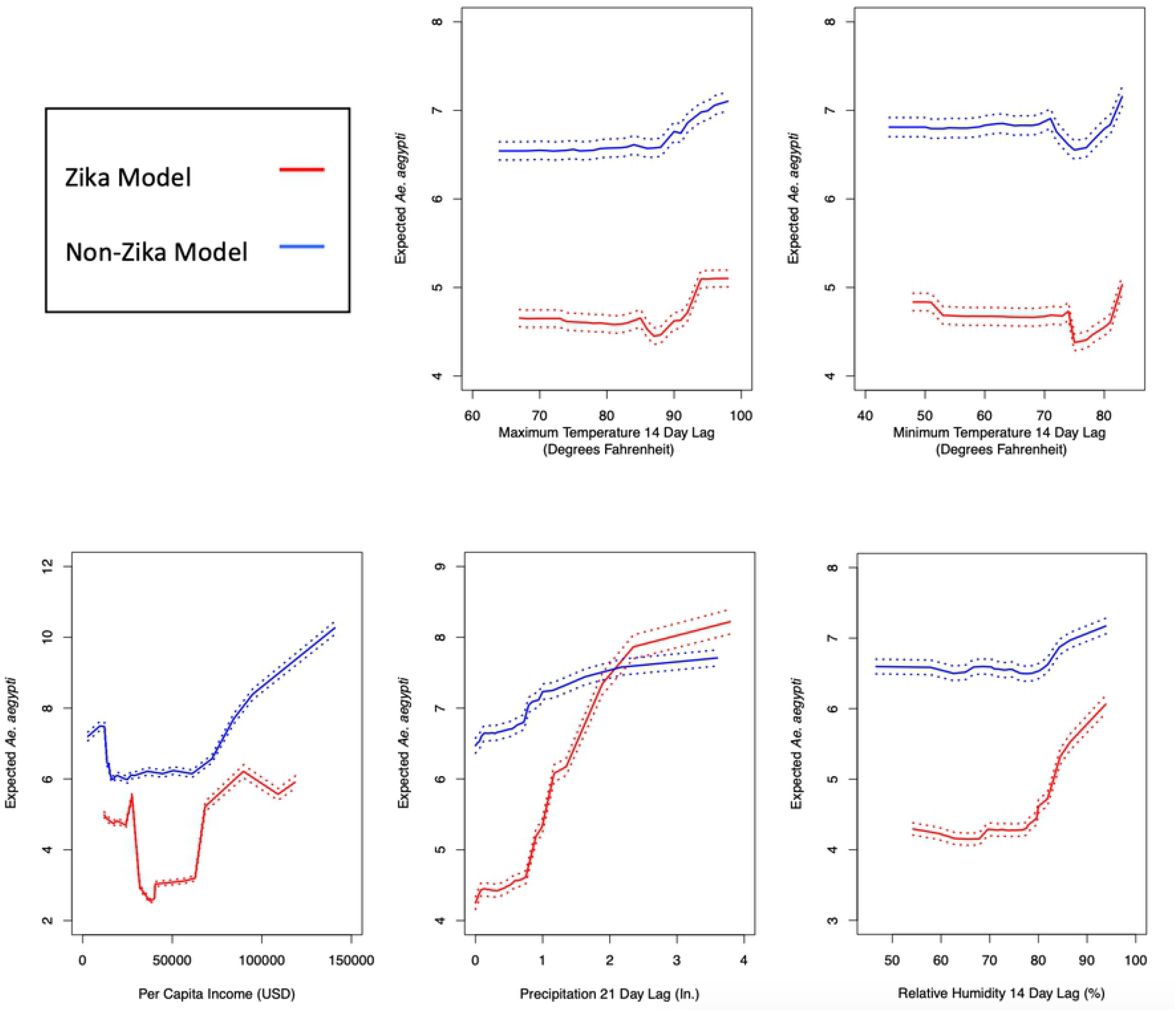
Partial plots of select variables in Random Forest models. These plots were generated for trap counts in areas affected by Zika during the 2016 outbreak (red) and unaffected areas. Dashed lines represent ±2 standard errors from expected *Aedes aegypti* counts per trap.

By cross-predicting the data, differences in the models are shown (Fig 6). Attempting to predict the Zika counts with the non-Zika model results in a weaker prediction than with the Zika model (RMSE = 11.74 > RMSE = 7.52). Similarly, attempting to predict the non-Zika counts with the Zika model results in a weaker prediction than with the non-Zika model (RMSE = 10.16 > RMSE = 5.73). Differences in model predictions are further shown below (Table 4) using a set of predictors.

**Fig 6.**
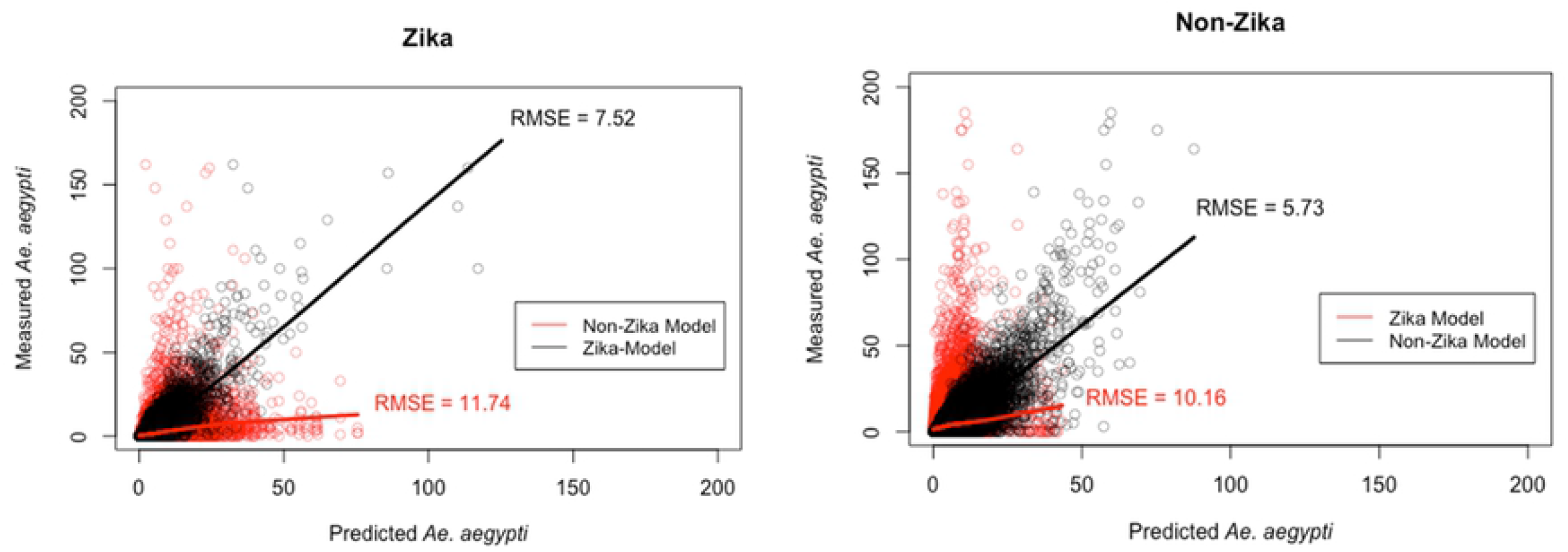
Predicted versus measured values for *Aedes aegypti* trap counts. These plots are shown for regions affected by local transmission of Zika during the 2016 outbreak (left) and unaffected regions (right), as predicted with random forest model trained on data from the respective region’s traps (black) or the opposite region’s traps (red).

**Table 4.**
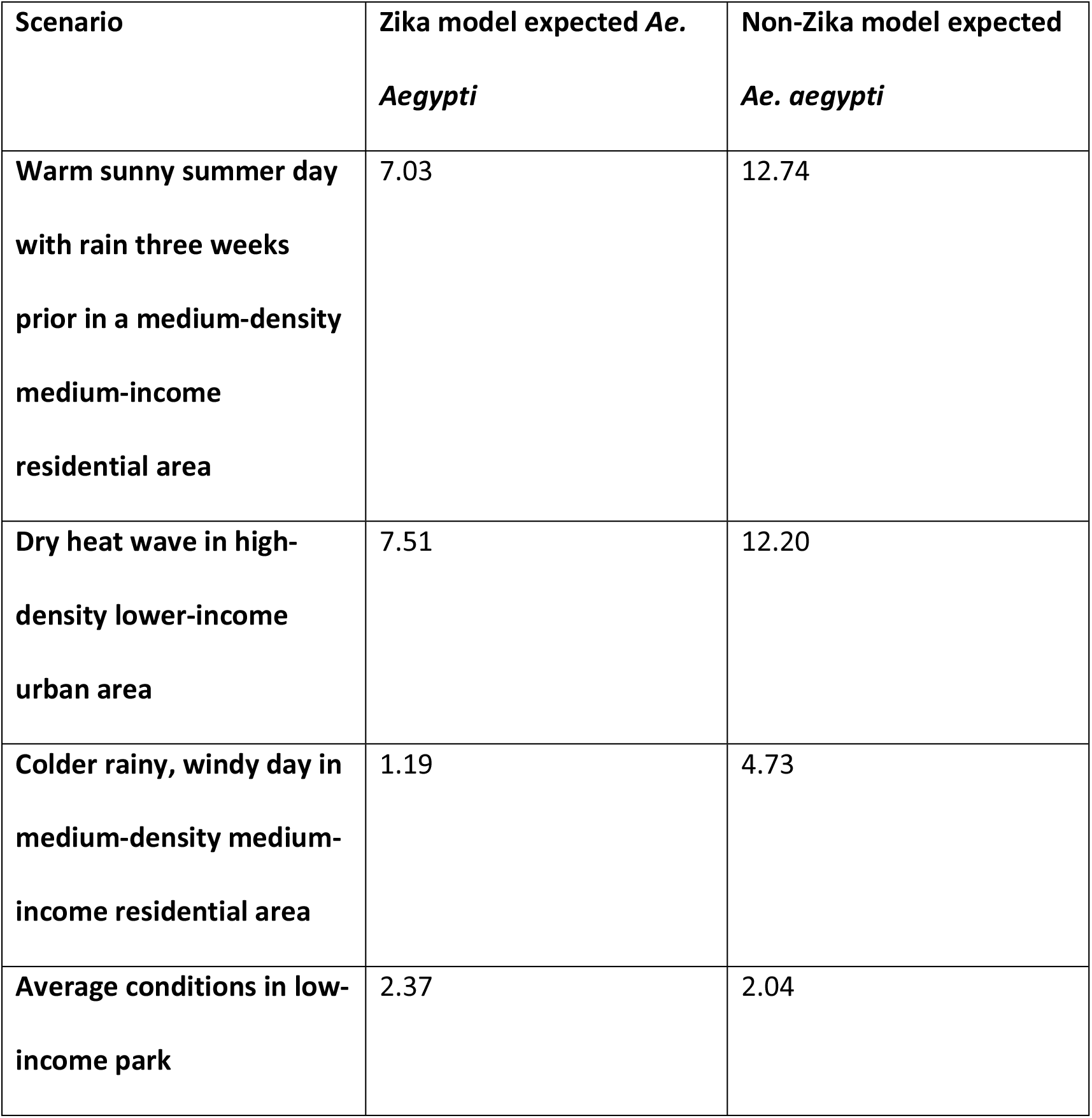
Sample predictions for given model input conditions. Predictions were made using random forest models trained on data from traps located in regions affected by local transmission of Zika during the 2016 outbreak (“Zika model”) and unaffected regions (“Non-Zika model”). A full list of input parameters is included in the Methods section.

## Discussion

The two hypotheses for this study were supported by the analysis. Temporally, while there is significant seasonal variation in the populations, the significant autocorrelation out to long-lengths, approximately half of a year or greater, in both regions implies that the populations do not rapidly fluctuate. However, we are unable to understand the variation in *Ae. aegypti* populations on a daily basis because of the nature of weekly collections. As weather conditions, especially precipitation, vary on a weekly basis, lacking daily data makes it hard to visualize the true temporal fluctuation in the data. More generally, it is possible that faster fluctuating vector populations may provide a greater risk for disease outbreaks, though this claim is not justified with prior studies. Spatially, there is significant variation in the number of *Ae. aegypti* collected per trap across different regions of the county. The traps with the highest average counts are only located in the more urbanized areas of the county (Fig 1). Even among nearby traps in these areas however, average trap count can vary greatly between traps in fairly close geographic proximity, indicating the major effects of the urban microgeography on *Ae. aegypti* populations.

The second hypothesis is also supported by this analysis. Random Forest model results suggest that long-term weather variables are more important determining factors of mosquito populations in areas affected by Zika whereas day-of-collection weather conditions and land-use variables are more important determining factors of mosquito populations in unaffected areas (Fig 4). Minimum daily temperature, maximum daily temperature, relative humidity two weeks before collection, and precipitation three weeks before collection were all more effective predictors of trap counts in areas affected by Zika than in unaffected areas. Note from the partial plots (Fig 5) that not only were relative humidity and precipitation at longer lag times better predictors of *Ae. aegypti* trap counts, but they also had a larger effect on *Ae. aegypti* populations, as expected *Aedes aegypti* varied 50-100% over the relative humidity and precipitation lagged conditions in Zika regions and only about 20% in non-Zika regions. All three of the land-use values were much better predictors of *Ae. aegypti* populations in non-Zika regions than in Zika regions, suggesting that the local urban landscape is more important to determining these populations. Day-of-collection weather conditions may affect mosquito activity levels and trap strength, but they are not, in the majority of cases, predictive of or affecting the true vector mosquito population. Nonetheless, these conditions are more important determining factors in the number of mosquitoes trapped in non-Zika regions than Zika-regions.

There are limitations to this study. Firstly, mosquito collections began in response to the Zika outbreak, so it is not possible to compare populations during the initial outbreak. As more traps were placed over the four-year period, trap quantity, spacing, and time frame were not constant between both regions. Simultaneously, mosquito control efforts around the county have not been controlled for in this study. Insecticide spraying, drainage of standing water, and other actions were taken by Miami-Dade County over the study period. These are factors that may contribute to the lower R^2^ values of the random forest models (Table 3). Additionally, they may contribute to populations in areas affected by Zika being generally lower than populations in unaffected areas as mosquito control efforts were focused more strongly on areas affected by local transmission of Zika[3] (Table 2). However, the random forest regression provides a method to understand the partial effects of relevant weather and spatial variables. Thus, while the overall models may be made weaker by greater variance in the data, the effects of the weather and spatial variables should be able to be inferred despite not accounting for population control efforts.

Other limitations of the study are a lack of trap-specific data on weather variables. All weather conditions may vary over the county. Weather conditions at each trap were not measured and therefore cannot be included in the analysis. This lack of data would only skew the results severely in the case that weather conditions vary differently over the Zika and non-Zika regions, but it is expected that both regions have similar variations in weather conditions. Temporal autocorrelation between weather variables at various lag times used is not too large for most variables but is necessary to mention for daily minimum and maximum temperature. For the entire prediction, the random forest model is not biased by collinearity between variables, as it will separate predictive strength between the variables in question. However, VIMP scores and partial plots may be affected for correlated variables as the model is forced to choose between variable importance and variable effect between similar variables. The collinearity between temperature measurements does not appear to have noticeably altered either the VIMP scores or the partial plots of any of the random forest runs.

For Miami-Dade County Mosquito Control, these results provide evidence that the current approach to *Aedes aegypti* population control has been sufficient. Populations in each region have been gradually declining, especially during peak season, the time period most important for disease transmission. A regional approach to insecticide and larvicide spraying is justified when disease transmission is likely and *Ae. aegypti* populations are exceedingly high. The exact population thresholds will vary based on human populations and disease transmission efficiency and will need to be determined by individual modeling efforts. The individual result of this study, that rainfall at about three weeks lag time appears to be a more significant determining factor of rainfall in some regions than others, indicates that the “drain and cover” motto should target these regions especially. It should also target areas with lower average per capita incomes and residential areas. Drainage should occur as soon as possible after any large rainfall event (> 0.8 inches).

In future studies, it may be valuable to utilize more variables including vegetation at the trap site, human population density in the surrounding region, and populations of competing mosquito species which may all provide additional predictive strength. This analysis could be repeated in other cities on a neighborhood basis to understand which areas may be most susceptible to outbreaks from vector-borne diseases. More simply, it might also be mimicked with other disease-vector mosquito species in Miami-Dade County to observe if similar patterns hold, especially as the threat of other vector-borne diseases like Dengue virus and West Nile Virus remains.

These results show the complexity of mosquito populations in urban environments. Microgeographic conditions are determining factors throughout the entire life cycle of disease-vector species and dictate important spatiotemporal variance in their populations. In turn, spatiotemporal variability in these populations is the foundation for the spread of vector-borne diseases. These nuances, outlined here in this study and others, should guide the efforts of epidemiologists and vector control experts in modeling and reducing vector-species populations. As climate change continues to affect global weather patterns, vector-species populations are expected to shift their ranges and behaviors. As this change occurs, microgeographic analysis like that conducted here can play a critical role in protecting public health.

## Acknowledgments

Thanks are given to the Miami-Dade County Mosquito Control who provided raw data for this study. The MDCMC performed the trap placement, trap collection, mosquito sexing, and mosquito species identification that was necessary for this study.

## Supporting information

**S1 Table.**
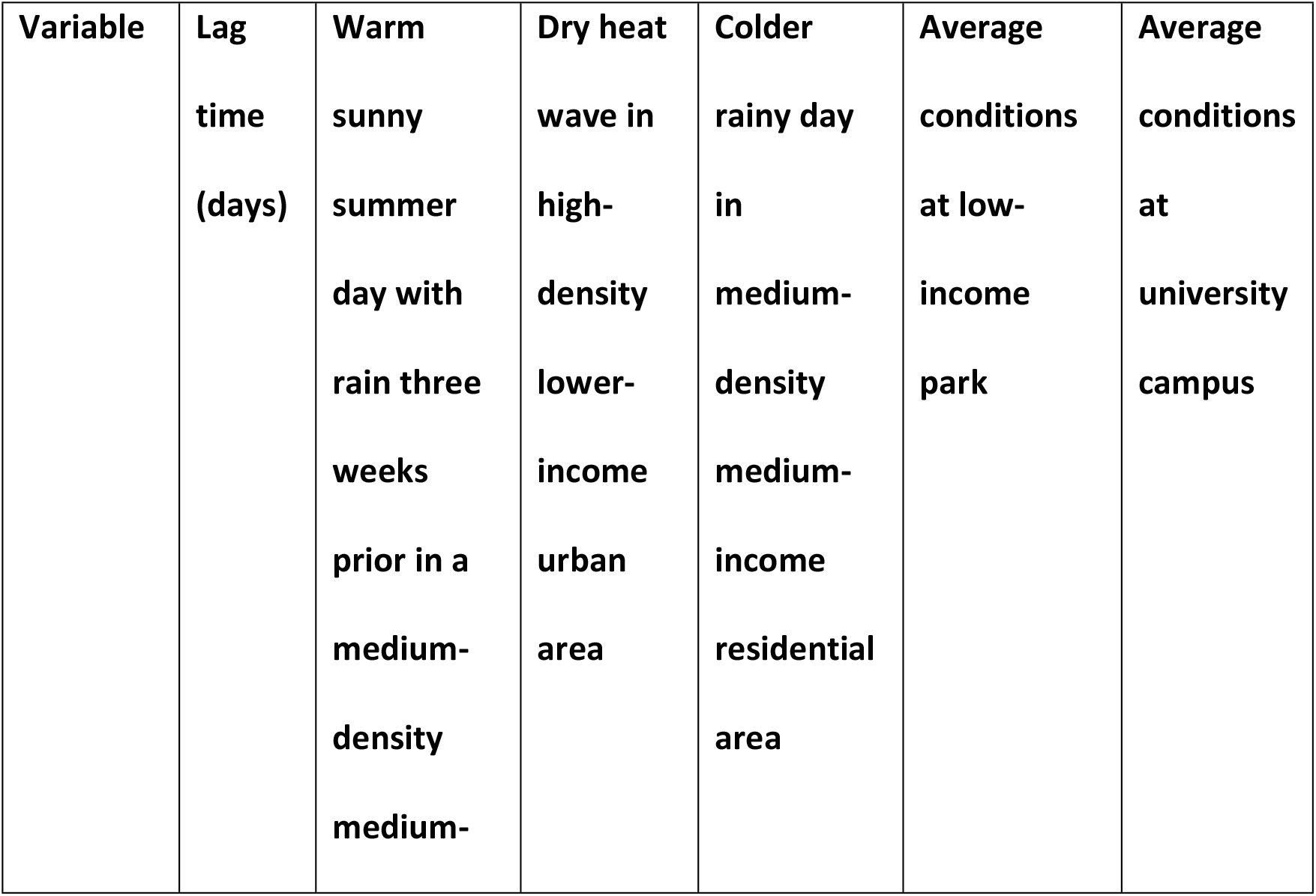
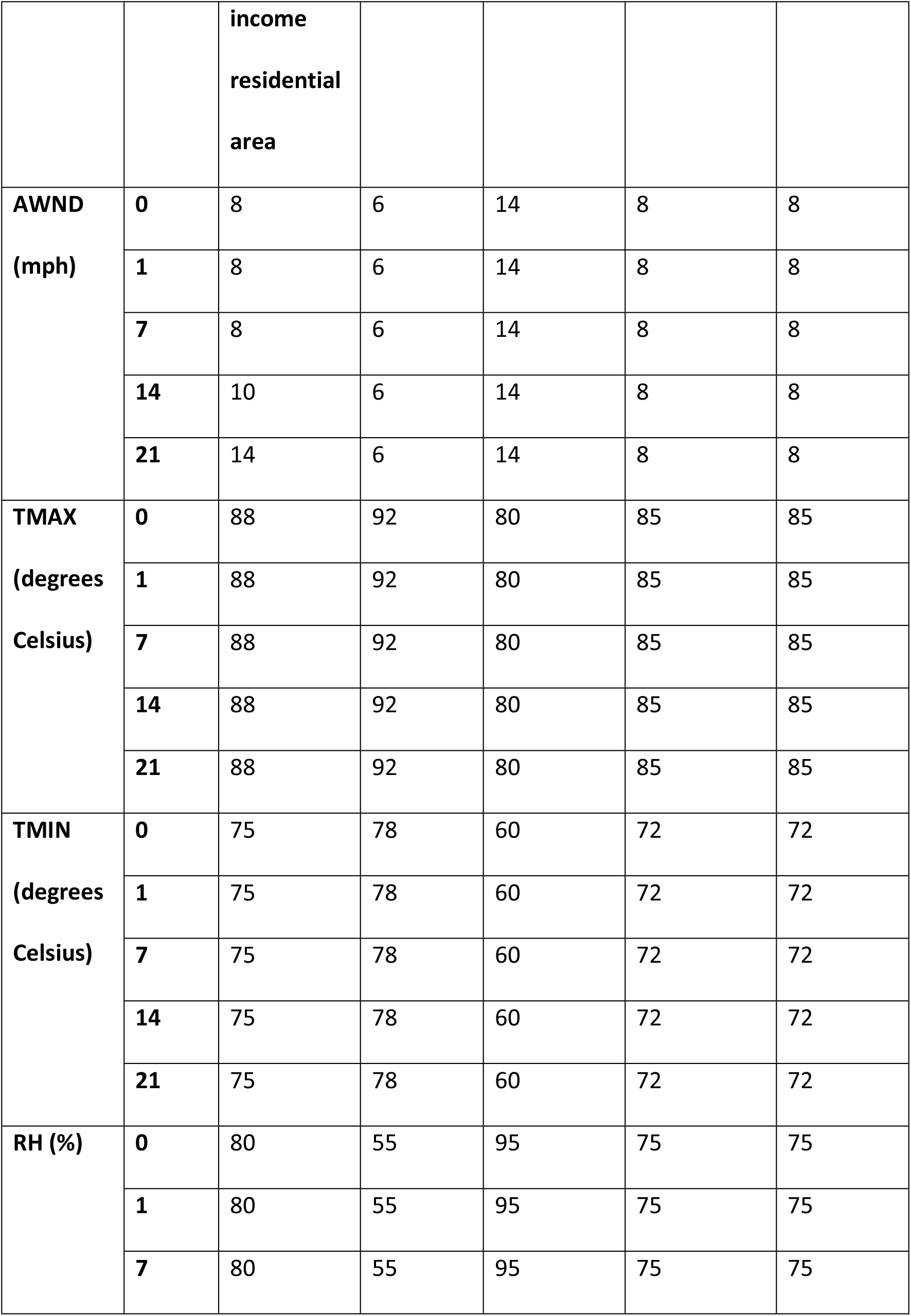
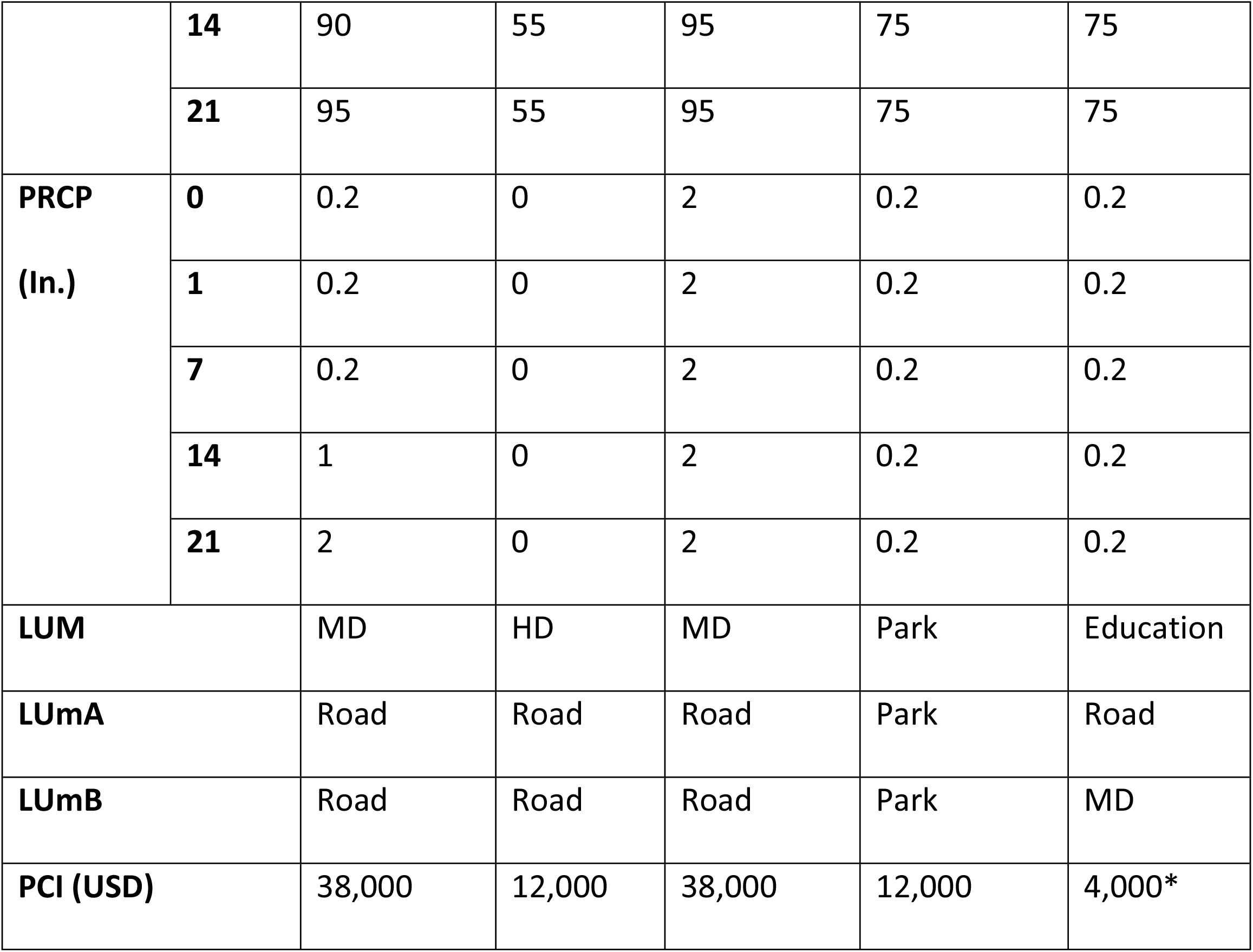
Variable values for sample predictions. Predictions are given in Table 4. Variable abbreviations can be found in Table 1. *Note that per capita income measured on university campuses are generally low as they capture student incomes.

## References

1. Wilke ABB, Vasquez C, Medina J, Carvajal A, Petrie W, Beier JC. Community Composition and Year-round Abundance of Vector Species of Mosquitoes make Miami-Dade County, Florida a Receptive Gateway for Arbovirus entry to the United States. Scientific Reports. 2019;9(1). doi: 10.1038/s41598-019-45337-2.

2. Likos A, Griffin I, Bingham AM, Stanek D, Fischer M, White S, et al. Local Mosquito-Borne Transmission of Zika Virus - Miami-Dade and Broward Counties, Florida, June-August 2016. Morbidity and Mortality Weekly Report. 2016;65(38):7. doi: 10.2307/24858999.

3. McAllister JC, Porcelli M, Medina JM, Delorey MJ, Connelly CR, Godsey MS, et al. Mosquito Control Activities during Local Transmission of Zika Virus, Miami-Dade County, Florida, USA, 2016. Emerg Infect Dis. 2020;26(5):881–90. doi: 10.3201/eid2605.191606. PubMed PMID: 32310079.

4. Petersen LR, Jamieson DJ, Powers AM, Honein MA. Zika Virus. N Engl J Med. 2016;374(16):1552–63. Epub 2016/03/31. doi: 10.1056/NEJMra1602113. PubMed PMID: 27028561.

5. Rocklöv J, Dubrow R. Climate change: an enduring challenge for vector-borne disease prevention and control. Nature Immunology. 2020;21(5):479–83. doi: 10.1038/s41590-020-0648-y.

6. Ogden NH, Lindsay LR, Ludwig A, Morse AP, Zheng H, Zhu H. Weather-based forecasting of mosquito-borne disease outbreaks in Canada. Can Commun Dis Rep. 2019;45(5):127–32. Epub 2019/07/10. doi: 10.14745/ccdr.v45i05a03. PubMed PMID: 31285703; PubMed Central PMCID: PMCPMC6587691.

7. Smith MW, Willis T, Alfieri L, James WHM, Trigg MA, Yamazaki D, et al. Incorporating hydrology into climate suitability models changes projections of malaria transmission in Africa. Nature Communications. 2020;11(1). doi: 10.1038/s41467-020-18239-5.

8. Wilke ABB, Medeiros-Sousa AR, Ceretti-Junior W, Marrelli MT. Mosquito populations dynamics associated with climate variations. Acta Tropica. 2017;166:343–50. doi: 10.1016/j.actatropica.2016.10.025.

9. Karki S, Brown WM, Uelmen J, Ruiz MOH, Smith RL. The drivers of West Nile virus human illness in the Chicago, Illinois, USA area: Fine scale dynamic effects of weather, mosquito infection, social, and biological conditions. PLOS ONE. 2020;15(5):e0227160. doi: 10.1371/journal.pone.0227160.

10. Ezanno P, Aubry-Kientz M, Arnoux S, Cailly P, L’Ambert G, Toty C, et al. A generic weather-driven model to predict mosquito population dynamics applied to species of Anopheles, Culex and Aedes genera of southern France. 2015;120(1):39–50. doi: 10.1016/j.prevetmed.2014.12.018.

11. Caldwell JM, Labeaud AD, Lambin EF, Stewart-Ibarra AM, Ndenga BA, Mutuku FM, et al. Climate explains geographic and temporal variation in mosquito-borne disease dynamics on two continents. 2020.

12. Ayanlade A, Nwayor IJ, Sergi C, Ayanlade OS, Di Carlo P, Jeje OD, et al. Early warning climate indices for malaria and meningitis in tropical ecological zones. Scientific Reports. 2020;10(1). doi: 10.1038/s41598-020-71094-8.

13. Robert MA, Christofferson RC, Silva NJB, Vasquez C, Mores CN, Wearing HJ. Modeling Mosquito-Borne Disease Spread in U.S. Urbanized Areas: The Case of Dengue in Miami. PLOS ONE. 2016;11(8):e0161365. doi: 10.1371/journal.pone.0161365.

14. Rose NH, Sylla M, Badolo A, Lutomiah J, Ayala D, Aribodor OB, et al. Climate and Urbanization Drive Mosquito Preference for Humans. Current Biology. 2020. doi: 10.1016/j.cub.2020.06.092.

15. Wilke ABB, Caban-Martinez AJ, Ajelli M, Vasquez C, Petrie W, Beier JC. Mosquito Adaptation to the Extreme Habitats of Urban Construction Sites. Trends in Parasitology. 2019;35(8):607–14. doi: 10.1016/j.pt.2019.05.009.

16. Wilke ABB, Vasquez C, Petrie W, Beier JC. Tire shops in Miami-Dade County, Florida are important producers of vector mosquitoes. PLOS ONE. 2019;14(5):e0217177. doi: 10.1371/journal.pone.0217177.

17. Wilke ABB, Vasquez C, Carvajal A, Moreno M, Diaz Y, Belledent T, et al. Cemeteries in Miami-Dade County, Florida are important areas to be targeted in mosquito management and control efforts. PLOS ONE. 2020;15(3):e0230748. doi: 10.1371/journal.pone.0230748.

18. Wilke ABB, Carvajal A, Vasquez C, Petrie WD, Beier JC. Urban farms in Miami-Dade county, Florida have favorable environments for vector mosquitoes. PLOS ONE. 2020;15(4):e0230825. doi: 10.1371/journal.pone.0230825.

19. Wilke ABB, Vasquez C, Mauriello PJ, Beier JC. Ornamental bromeliads of Miami-Dade County, Florida are important breeding sites for Aedes aegypti (Diptera: Culicidae). Parasites & Vectors. 2018;11(1):283. doi: 10.1186/s13071-018-2866-9.

20. Wilke ABB, Chase C, Vasquez C, Carvajal A, Medina J, Petrie WD, et al. Urbanization creates diverse aquatic habitats for immature mosquitoes in urban areas. Scientific Reports. 2019;9(1). doi: 10.1038/s41598-019-51787-5.

21. Moise IK, Riegel C, Muturi EJ. Environmental and social-demographic predictors of the southern house mosquito Culex quinquefasciatus in New Orleans, Louisiana. Parasites & Vectors. 2018;11(1). doi: 10.1186/s13071-018-2833-5.

22. Segal M, Xiao Y. Multivariate random forests. WIREs Data Mining and Knowledge Discovery. 2011;1(1):80–7. doi: https://doi.org/10.1002/widm.12.

23. Ong J, Liu X, Rajarethinam J, Kok SY, Liang S, Tang CS, et al. Mapping dengue risk in Singapore using Random Forest. PLOS Neglected Tropical Diseases. 2018;12(6):e0006587. doi: 10.1371/journal.pntd.0006587.

24. Mussumeci E, Codeço Coelho F. Large-scale multivariate forecasting models for Dengue - LSTM versus random forest regression. Spatial and Spatio-temporal Epidemiology. 2020;35:100372. doi: https://doi.org/10.1016/j.sste.2020.100372.

25. Georganos S, Grippa T, Niang Gadiaga A, Linard C, Lennert M, Vanhuysse S, et al. Geographical random forests: a spatial extension of the random forest algorithm to address spatial heterogeneity in remote sensing and population modelling. Geocarto International. 2021;36(2):121–36. doi: 10.1080/10106049.2019.1595177.

26. Wilke ABB, Carvajal A, Medina J, Anderson M, Nieves VJ, Ramirez M, et al. Assessment of the effectiveness of BG-Sentinel traps baited with CO2 and BG-Lure for the surveillance of vector mosquitoes in Miami-Dade County, Florida. PLOS ONE. 2019;14(2):e0212688. doi: 10.1371/journal.pone.0212688.

27. Placement and effects of Biogents mosquito traps. https://usbiogentscom/wp-content/uploads/Placement-and-effect-of-Biogents-mosquito-traps-enpdf. Biogents USA Website: Biogents.

28. National Online Weather Database NOAA National Centers for Environmental Information: National Oceanic and Atmospheric Administration; [cited 2020 June 25, 2020]. Available from: https://www.ncdc.noaa.gov/cdo-web/datasets/GHCND/stations/GHCND:USW00012839/detail.

29. U.S. Local Climatological Data (LCD). In: National Centers for Environmental Information N, NOAA, U.S. Department of Commerce, National Weather Service N, U.S. Department of Commerce, U.S. Air Force USDoD, Federal Aviation Agency USDoT, editors. National Centers for Environmental Information, NESDIS, NOAA, U.S. Department of Commerce.

30. Miami-Dade County. Miami-Dade County Open Data Hub 2021 [updated October 21, 2021; cited 2021 September 27, 2021]. Available from: https://gis-mdc.opendata.arcgis.com/.

31. ArcGIS [GIS Software]. 10.8.1 ed: Environmental Systems Research Institute, Inc.

32. Per Capita Income in the Past 12 Months (In 2019 Inflation-Adjusted Dollars): United States Census Bureau; [cited 2021 October 1, 2021]. Available from: https://data.census.gov/cedsci/table?q=per%20capita%20income%20by%20census%20tract&tid=ACSDT1Y2019.B19301.

33. Per Capita Income: Miami Matters; [cited 2021 October 1, 2021]. Available from: http://www.miamidadematters.org/indicators/index/view?indicatorId=15&localeId=138823.

34. Venables WN, Ripley BD. Modern Applied Statistics with S (MASS). 7.3-54 ed 2021.

35. R Core Team. R: A Language and Environment for Statistical Computing. 4.11 ed: R Foundation for Statistical Computing; 2021.

36. RStudio Team. RStudio: Integrated Development for R. RStudio, PBC; 2020.

37. Microsoft Corporation. Microsoft Excel for Mac. 16.54 ed 2021.

38. Ishwaran H, Kogalur UB. Fast Unified Random Forests for Survival, Regression, and Classification (RF-SRC). 2.12.1 ed 2021.

